# Relating Monosynaptic and Functional Connectivity - Complementary Perspectives on Neural Computation -

**DOI:** 10.1101/2025.05.05.652327

**Authors:** Shigeru Shinomoto, Yasuhiro Tsubo

**Affiliations:** Graduate School of Biostudies, Kyoto University, Kyoto 606-8501, Japan; Research Organization of Open Innovation and Collaboration, Ritsumeikan University, Osaka 567-8570, Japan; College of Information Science and Engineering, Ritsumeikan University, Osaka 567-8570, Japan

## Abstract

Information processing in the brain is thought to result from the coordination of large-scale neuronal activity, but this occurs on a fine neural circuit. Here, we investigate their relationship. First, we estimate monosynaptic connectivity by applying an advanced analysis method to spike trains recorded with high-density microelectrodes and confirm that the estimated neuronal wiring is largely consistent with neuroanatomical and neurophysiological evidence. Second, we simulate calcium imaging signals from the same dataset and confirm that the estimated functional connectivity is influenced by shared inputs and population synchronization on slower timescales. Notably, even with unrealistically fast calcium dynamics, the functional connectivity is only partially consistent with the monosynaptic connectivity. These findings suggest the complementary roles for monosynaptic and functional connectivity: the former provides circuit-level specificity, while the latter reflects emergent system-wide patterns of activity. We propose that an integrative approach combining both perspectives is essential for understanding circuit-level computation in the brain.

## INTRODUCTION

With the development of large-scale recording techniques, there has been a surge of interest in capturing the communication that takes place between different parts of the brain. Researchers have proposed to detect functional connectivity by capturing significant synchronizations between their activity, with the hope of understanding how the brain supports cognition and behavior, or capturing brain dysfunction when it occurs [1–3]. Recently, two-photon microscopy technology is rapidly evolving to allow parallel recording of a large number of neurons, opening up the possibility of drawing a huge detailed map of neuronal functional connectivity [4–9]. Despite these advances, it is still poorly understood how such neuronal co-activity emerges on the underlying hard-wired neuronal circuitry.

It seems difficult for the two-photon microscopy technology to elucidate neuronal hardware wiring because calcium dynamics on the order of 100 ms are much slower than monosynaptic signal transmission between neurons, which occurs in a few ms. In contrast, microelectrode technology, which can determine neuronal firing with sub-millisecond accuracy, has the potential to reveal the causal antecedents and sequelae of neuronal activation. The number of recordable neurons is also increasing due to the development of high-density electrodes such as Neuropixels [10, 11], and analysis methods for estimating monosynaptic connections from parallel spike trains have been developed and implemented [12–29].

Here, we investigate how functional connectivity relates to the underlying neuronal wiring structure. To this end, we first estimate the monosynaptic connections between recorded neurons using a state-of-the-art analysis method. By analyzing a large number of spike trains recorded with high-density microelectrodes, we examine the consistency of the estimated monosynaptic connections with neuroanatomical and neurophysiological evidence. Second, we compute the functional connections between the same set of neurons by simulating two-photon imaging signals from the spike trains. To do this, we convolve a kernel representing a calcium influx for each spike and the subsequent pumping. By varying the timescale of the kernel, we investigate how well the functional connectivity can be correlated with the monosynaptic connectivity, and in particular, whether the functional connectivity can ultimately represent the monosynaptic connectivity in the limit of the shortest timescale. Third, we perform a simulation of an artificial neural network consisting of estimated monosynaptic connections to see how well the simulated network can reproduce the real functional connections. We also apply weak stimuli to a subset of neurons to see how functional connectivity can be affected by external stimuli.

## RESULTS

### Estimating monosynaptic connections

Given a set of neuronal spike trains, a state-of-the-art analysis method determines whether any pair of neurons are monosynaptically connected based solely on their spike timing information, without relying on any other knowledge such as the distance between the recorded neurons. This is a difficult task, as if we are trying to infer the relationship between any pair of people from their chat times alone, without using other cues such as their social community or where they live. Although conventional algorithms have made many incorrect estimates, subsequent innovations have greatly improved the accuracy of the estimates [16–26]. While efforts are still underway to improve and compare the performance of analysis methods [27, 28], we focus on evaluating the reliability of the latest analysis technique by itself, and summarizing the information about the neuronal circuitry specific to each brain region. Here, we adopt our latest algorithm ShinGLMCC [29] and analyze spike trains recorded with a single probe consisting of high-density electrodes, Neuropixels [10, 11].

### Neural circuits in different brain regions

Given a numerical data of recorded spike trains, the ShinGLMCC estimation algorithm provides a table of estimated connections for all neuron pairs, along with another table of their statistical significance *α* (Figure 1(a)). Through-out this paper, we have determined monosynaptic connections by thresholding each directed pair with *α* ≤ 0.001 (METHODS).

**FIG. 1.**
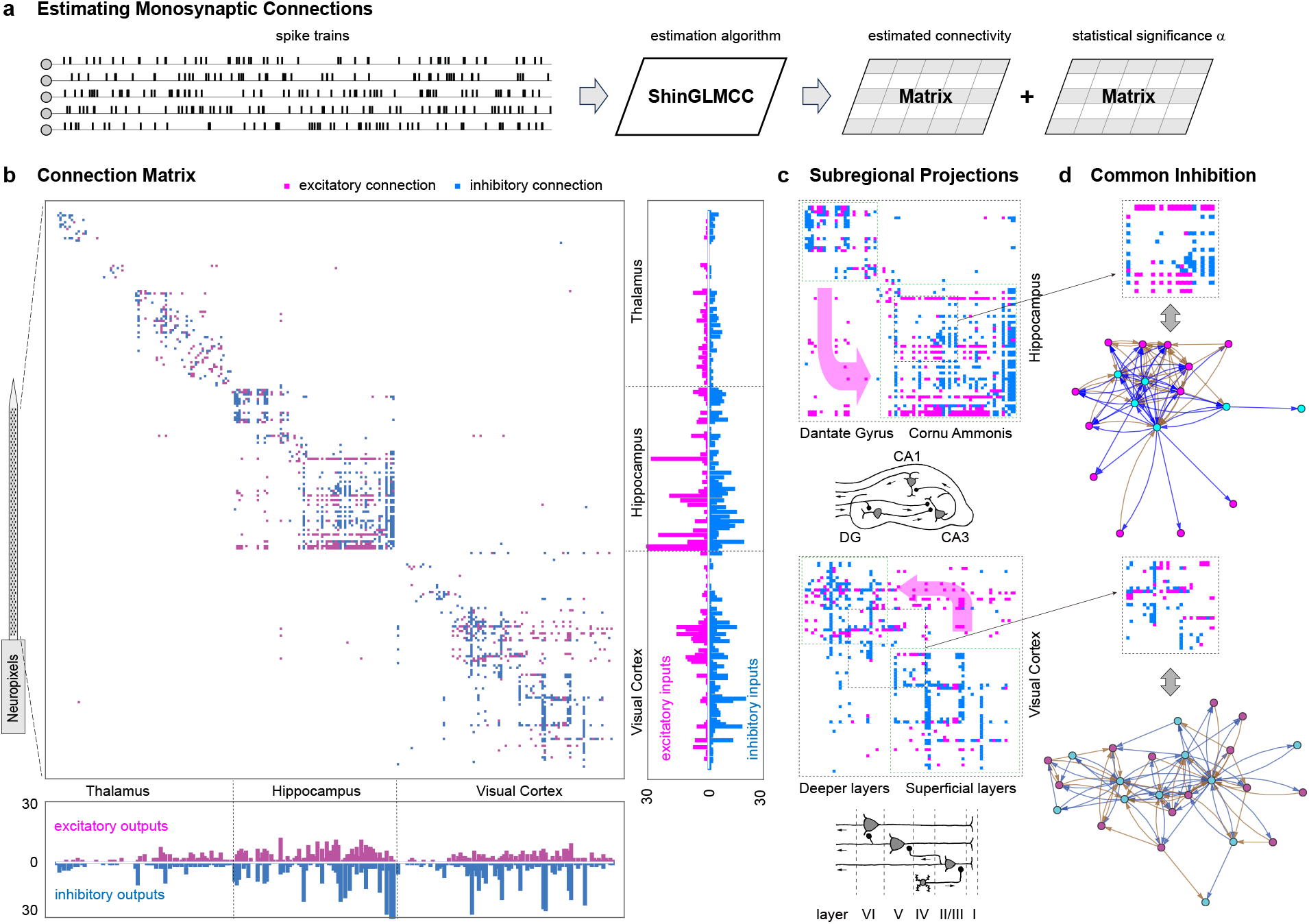
Connection matrix of a neuronal ensemble recorded with a single high-density electrode. (a) The procedure for estimating monosynaptic connections. (b) A complete matrix of 242 neurons arranged along the axial direction of an electrode, covering multiple regions including thalamus, hippocampus, and visual cortex (The dataset used here is publicly available from the original study [10]). Excitatory and inhibitory connections are colored in magenta and light blue, respectively. The horizontal and vertical axes represent output and input indices, respectively. The number of excitatory and inhibitory outputs for each neuron is shown below the matrix, while the number of excitatory and inhibitory inputs for each neuron is shown on the right. (c) Magnified view of the connections in the hippocampus and visual cortex. Significant unidirectional excitatory projections from the dentate gyrus (DG) to the cornu ammonis (CA) in the hippocampus and from supragranular layers to infragranular layers in the visual cortex are indicated by red arrows. The anatomically known subregional projections in each region are shown below. (d) Further magnification of parts of the hippocampus and visual cortex, showing reciprocal connections between inhibitory interneurons and excitatory pyramidal neurons. Network diagrams are drawn below each connection matrix. Putative inhibitory and excitatory neurons are represented by magenta and cyan circles, and the estimated inhibitory and excitatory connections are colored in brown and blue, respectively.

Figure 1(b) shows the connection matrix estimated by applying ShinGLMCC to spike trains recorded in parallel from a rat. In the matrix, presynaptic and postsynaptic neurons are arranged in order along the axial direction of a single probe of Neuropixels from left to right and top to bottom, respectively. Experimental researchers recorded electrical signals, sorted individual neuronal spikes, and also identified the possible spatial location of the recorded neurons. In the dataset we analyzed here, the total length of the probe spanned several brain regions consisting of the thalamus, hippocampus, and visual cortex[10, 11]. At the periphery of the matrix, we indicate the presumed boundaries of these brain regions.

While ShinGLMCC determines monosynaptic connections without reference to the recorded positions, the estimated connections turned out to be clearly well localized to each brain region. In addition, the patterns of neuronal connectivity differ significantly between different brain regions. Interneuronal connections in the thalamus are fewer and more localized than in the hippocampus and cortex, consistent with neuroanatomical and neurophysiological knowledge. In contrast, connections in the hippocampus and visual cortex are widespread throughout each region.

The numbers of outgoing excitatory and inhibitory connections for each neuron are shown below the matrix. As with cortical neurons, the output connections for each neuron are known to be uniquely polarized as either excitatory or inhibitory. Because the analysis is performed statistically on a finite number of spike events, any estimate is inevitably subject to error. Note that errors may be due not only to the estimation algorithm, but also to possible errors in the spike sorting of the experimental data.

From this histogram of outgoing connections, we can see that putatively inhibitory neurons tend to innervate a greater number of connections than putatively excitatory neurons. This tendency is particularly strong in the cornu ammonis (CA) region of the hippocampus. It is noteworthy that some inhibitory neurons innervate the majority of recorded neurons in the CA region. Since the recorded neurons may represent only a small fraction of the total ensemble, it is likely that inhibitory interneurons in the CA region are connected to a very large number of neurons. The number of excitatory and inhibitory inputs to each neuron is shown on the right side of the matrix. The histogram shows that inhibitory neurons tend to have many excitatory inputs. This is consistent with the physiological knowledge that inhibitory interneurons tend to have reciprocal excitatory connections from the pyramidal neurons that they inhibit.

Figure 1(c) magnifies the hippocampus and a part of the visual cortex to focus on their individual intersubregional projections. There are prominent excitatory projections from the dantate gyrus (DG) to the CA in the hippocampus and from superficial layers to deeper layers in the visual cortex. These unidirectional projections are consistent with neuroanatomical knowledge [30–32].

Figure 1(d) depicts parts of the hippocampus and visual cortex to focus on the reciprocal connections between inhibitory interneurons and excitatory pyramidal neurons, as we have noted above. This circuit structure supports the idea of a common inhibition mechanism in which a “winner-takes-all” mechanism operates [33–35]. In the hippocampus, when a rat is at some location, place cells specific to that location activate inhibitory cells to suppress other place cells. When the rat moves to another location, other place cells activate the same inhibitory cells to suppress others. This is consistent with the knowledge that inhibitory cells are not place selective. In the visual cortex, inhibitory neurons also have reciprocal connections between many excitatory cells, but the number of connections is not as large as in the hippocampus [36].

### Estimating functional connections

Next, we investigate the relationship between functional and monosynaptic connectivity for the same set of neurons. To do this, we artificially generate two-photon signals from the spike trains that we used to estimate monosynaptic connections. The fluorescence signal represents the calcium concentration in each neuron; when it fires, calcium immediately enters the cell and is then slowly pumped out. We represent this process by convolving an exponential kernel function at each spike time (Figure 2(a)).

**FIG. 2.**
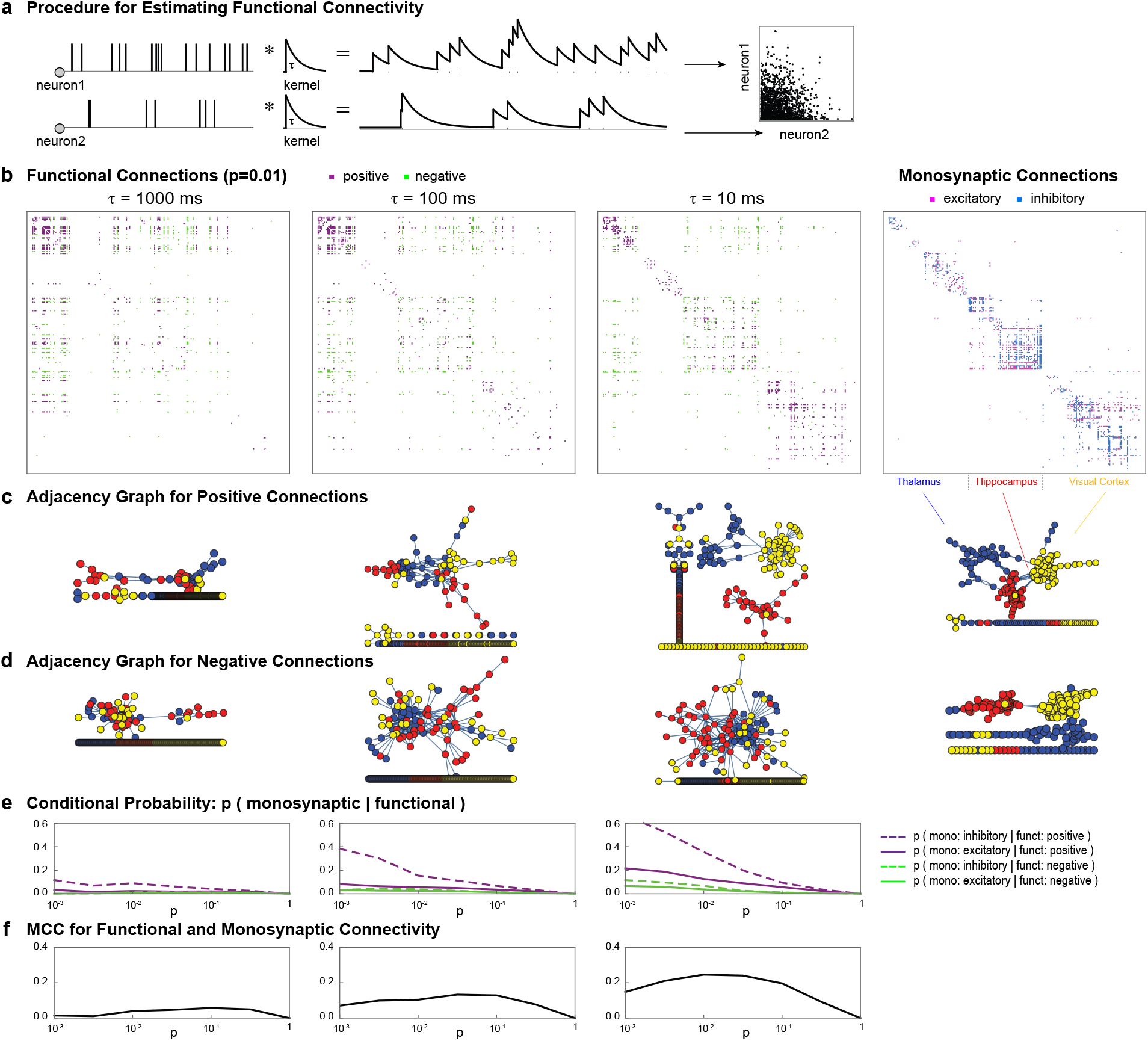
Functional and monosynaptic connections. (a) The procedure for artificially generating two-photon Ca signals from recorded spike trains, and computing functional connectivity from the Pearson correlation coefficient. (b) Functional and monosynaptic connectivity estimated for the same set of neurons recorded from the thalamus, hippocampus, and visual cortex. The timescale of a convolution kernel is chosen as *τ* = 1000 ms, 100 ms, or 10 ms. Positive and negative functional connections are colored purple and green, respectively. (c) and (d) Adjacency graphs for the positive and negative functional connections, with reference to the adjacency graphs for excitatory and inhibitory monosynaptic connections. Neurons of the thalamus, hippocampus, and visual cortex are colored blue, red, and yellow, respectively. (e) Conditional probability that a pair of neurons is monosynaptically connected, given that they are functionally connected. (f) Association between the presence or absence of functional and monosynaptic connectivity, represented by the Matthews correlation coefficient (MCC).

Given a set of time series of fluorescence signals, we compute functional connectivity using the standard method, namely, we compute the Pearson correlation coefficient between the time series for each pair of neurons, and then select the top *p* fraction of pairs that have the highest correlation among all pairs. A high correlation implies that the pairs of neurons are working together. We also look at the pairs that have the lowest correlation, which we call negative functional connectivity to distinguish it from positive one.

### Comparison of functional and monosynaptic connectivity

Figure 2(b) compares the functional and monosynaptic connections estimated for the neurons recorded from the thalamus, hippocampus, and visual cortex. Because the fluorescence signals are artificially generated, we can examine how the functional connections vary with the timescale of the kernel. Here we varied the time scale of a convolving kernel from *τ* = 1000 ms, 100 ms, to 10 ms. First, we observe that the positive functional connections are relatively more localized in three brain regions than the negative functional connections. Second, the positive functional connections become more localized as the time scale *τ* becomes shorter.

Figures 2(c) and (d) show the adjacency graphs of the positive and negative functional connections, with the nodes of three brain regions colored differently. Neurons of three regions are relatively well clustered with positive functional connections when the timescale is unrealistically short, *τ* = 10 ms. However, the clustering features are still not as clear as those for excitatory and inhibitory monosynaptic connections estimated by ShinGLMCC, as shown on the right. We tested an even shorter time scale *τ* = 1 ms, but the situation is similar and the functional connections do not approach the monosynaptic connections.

Because neuronal function can occur on a given circuit, it is naturally expected that neuronal co-activation, or functional connectivity, will occur for a pair of neurons that are monosynaptically connected. To see their relationship, we computed the conditional probability that each pair of neurons is monosynaptically connected, given that they are functionally connected (Figure 2(e)). However, the conditional probability turned out to be low on the realistic timescale of calcium dynamics of about *τ* = 1000 ms. On the timescale of 100 ms, which may be the shortest possible timescale in reality, the conditional probability of an inhibitory connection can be as high as 0.4 for pairs with very strong top *p* = 0.001 functional connections. In other words, a pair of neurons showing strong coactivation could be monosynaptically connected, but more likely have an inhibitory connection. Symbolic properties of monosynaptic and functional connectivity are summarized in the Table I.

Because we are making artificial time series, we can go to even shorter timescales, such as *τ* = 10 ms, which does not seem feasible with current technology. Then the conditional probabilities get higher. For example, pairs with highly selective *p* = 0.001 positive functional connectivity are likely to be monosynaptically connected. It is also true that the probability of an inhibitory connection is higher than that of an excitatory connection. However, this does not mean that the functional connections are similar to the monosynaptic connections, because the conditional probabilities become lower for the weaker functional connectivity such as *p* = 0.01 or 0.1.

We estimate an association between functional and monosynaptic connectivity in terms of the Matthews correlation coefficient (MCC), which measures the balance of binary variables by taking into account their true and false positives and negatives [37]. Here we simplify the classifications into the presence or absence of connections, ignoring their signs, such as positive or negative functional connections and excitatory or inhibitory monosynaptic connections. The MCC varies with the probability *p* of selecting functional connections (Figure 2(f)). For high probabilities such as *p >* 0.1, the MCC is low because the less selective functional connectivity may not be associated with monosynaptic connectivity. On the contrary, for low probabilities such as *p* = 0.001, the pairs with strong functional connectivity tend to have monosynaptic connectivity, but the other majority lose the association. As a result, the MCC peaks between *p* = 0.01 and *p* = 0.1. The peak value of the MCC increases as the timescale is decreased from *τ* = 1000 ms to *τ* = 10 ms, indicating that the functional connectivity is closer to the monosynaptic connectivity. Note, however, that the value is still much less than the perfect match, or MCC = 1, indicating that functional connectivity cannot represent monosynaptic connectivity even at the short timescale of *τ* = 10 ms. We also tested the shortest timescale of *τ* = 1 ms, but the situation did not change significantly. Thus, the functional connectivity does not represent the monosynaptic connectivity even in the limit of the shortest timescale.

### Numerical simulation of a network of estimated connections

While the recorded neurons are still a tiny fraction of the neurons in the brain, their monosynaptic connections exhibit specific features consistent with neuroanatomical knowledge. To investigate the extent to which the functional connections can be reproduced by the network of recorded neurons alone, here we perform a simulation of an artificial neural network consisting of the estimated connections.

A numerical simulation is performed using the spiking neuron model, the Multi-timescale Adaptive Threshold (MAT) model [38]. Synaptic connections between model neurons were provided by the connection matrix estimated from the real data set shown in Figure 1(b). In addition to these intranetwork connections, we added noise representing inputs from invisible neurons so that the firing rate of each neuron is equal to the recorded firing rate. This simulation has only one free parameter that multiplies the estimated monosynaptic connections to determine the strength of the intranetwork connectivity relative to the inputs from external invisible neurons. We chose this parameter so that the monosynaptic connections estimated from simulated spike trains become similar to the original ones (METHODS). As a result, these monosynaptic connection matrices showed good agreement, as can be seen by comparing the right ends of Figure 2(b) and Figure 3(a).

**FIG. 3.**
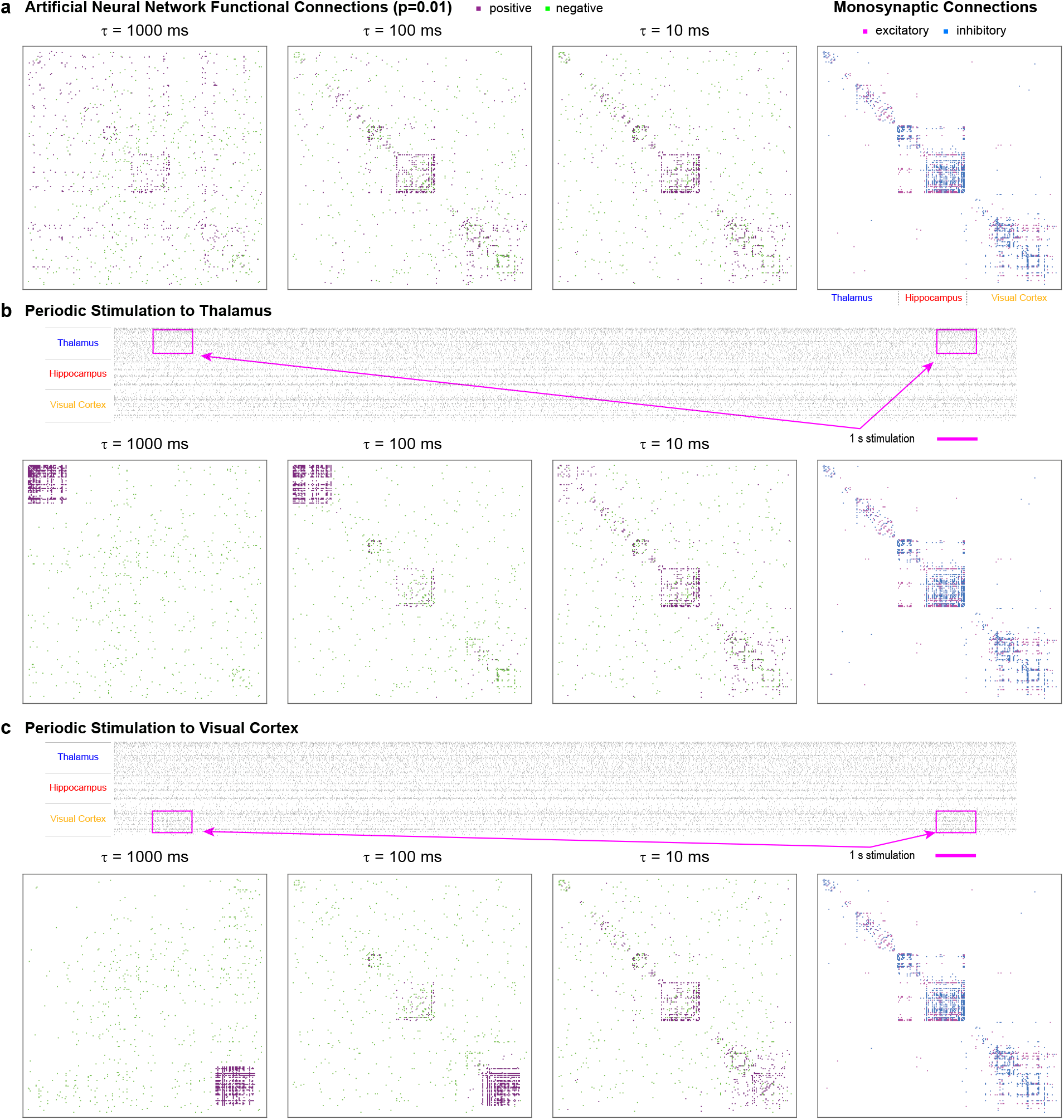
Functional and monosynaptic connections in an artificial neural network. (a) Connections estimated for spike trains generated by numerical simulation. The timescale of a convolution kernel is chosen to be *τ* = 1000 ms, 100 ms, or 10 ms. (b) and (c) Connections estimated for neural networks with weak stimulation applied to the thalamus and visual cortex, respectively. Raster diagrams show sample spike trains of neurons of even number indices for 23 seconds. The weak stimulus was applied repeatedly for 1 second every 20 seconds.

Three left panels of Figure 3(a) show the resulting functional connections for the timescales *τ* = 1000 ms, 100 ms, or 10 ms. Although these functional connections of an artificial neural network do not reproduce monosynaptic connections either, they are more similar to monosynaptic connectivity than those of the real spike trains shown in Figure 2(b), implying that the functional connectivity of the simulated spike trains tends to reflect the monosynaptic connectivity relatively more faithfully than the real spike trains. This may be because the simulation is performed with the isolated network, while the real network receives correlated inputs from external invisible neurons whose states change over time.

To see how the functional connectivity is affected by external stimulation, we ran simulations in which weak stimulation was applied to the thalamus or visual cortex. Namely, we repeatedly increased the excitatory noise to neurons in each area for 1 second in every 20 seconds (Figures 3(b) and (c)). The applied stimulus is quite modest, so that the firing rates of the stimulated neurons increased by about 1.5 times for the short period of time. Such a small perturbation has induced a very strong functional connectivity among these stimulated neurons, particularly for the case of a realistic timescale of *τ* = 1000 ms or 100 ms. This implies that the functional connectivity of the fluorescent calcium signal is sensitively influenced by common inputs to neurons.

On the contrary, the estimated monosynaptic connections were hardly affected by the stimulation, as evidenced by the MCCs between unstimulated and stimulated systems (right ends of Figures 3 (a), (b), and (c)) being greater than 0.98. This fact supports the robustness of the monosynaptic connection estimation method, ShinGLMCC.

## DISCUSSION

In the present study, we analyzed high-density recording data to estimate monosynaptic and functional interneuronal connectivity. First, we estimated direct neuronal wiring using the advanced analysis method. Second, we simulated calcium imaging signals and calculated functional connectivity. Third, we simulated a neural network consisting of the estimated connections. From these analyses, we learned that the state-of-the-art analysis method accurately captures the direct interaction between neurons by extracting the synaptic influence with a delay of a few ms, while the functional connectivity is strongly influenced by shared inputs, indirect interactions, and population synchronization. In particular, we found that functional connectivity at the time resolution of the calcium fluorescence signal is only partially consistent with the underlying monosynaptic connectivity. Even at unrealistically fast time resolutions, functional connectivity does not represent the interneuronal wiring.

We have shown that the latest monosynaptic estimation method, ShinGLMCC, not only makes inferences consistent with neuroanatomical and neurophysiological evidence, but also reveals new information about individual brain regions, such as the extensive innervation of inhibitory neurons in the hippocampus. A more important contribution of this estimation method may be its ability to provide concrete values for interneuronal connections, allowing realistic network simulations. These results are also due to the latest recording technology, which has made it possible to accurately monitor a large number of spike trains. Note that there still seem to be errors in spike sorting. We expect that the signal-to-noise ratio of the measurement and the sorting algorithm will continue to improve. We have confirmed that the latest connection estimation algorithm is a fairly reliable analysis tool, but we also expect that the algorithm to continue to improve.

Monosynaptic connectivity is estimated by judging whether the synaptic influence with a delay of a few ms is statistically significant in the cross-correlogram. This statistical inference requires a large number of spikes to determine the existence of a monosynaptic connection. Currently, we are using data from 1-2 hours of measurements. It would be desirable to have longer time measurements that would allow the estimation of even weaker connections. Note that the need for a long monitoring time is not a technical requirement of a specific analysis method, but essentially necessary to obtain statistical confidence from a finite number of spikes, as was given in equation (3) or Table 1 in the reference [24].

**TABLE 1.** Characterization of monosynaptic and functional connectivity.

|  | Monosynaptic | Functional |
| --- | --- | --- |
| Time scale | 1-5 ms | 100-1000 ms |
| Spatial scale | Local | Global |
| Role | Hardware | Software |
| Representation | Synaptic wiring | Neuronal dynamics |

Next, we estimated functional connectivity between the same set of neurons and examined its relationship to monosynaptic connectivity, because we expected that neuronal co-activation, or functional connectivity, would appear on top of the network of connected neurons and might correlate with monosynaptic connectivity. However, we found that functional connectivity at the time resolution of calcium fluorescence signals is a weak reflection of the underlying neuronal circuitry and does not approach monosynaptic connectivity even at the shortest time resolution.

Throughout this analysis, we have adopted the standard definition of functional connectivity as the co-activity of a pair of neurons, or the Pearson correlation of calcium dynamics. There are many variants of functional connectivity that aim to capture the directed flow of information in the brain, in particular by detecting the causal interaction or activation delay between nodes [39–41]. While these measures may add information about the dynamic aspect of the brain, it still seems difficult to relate them to the underlying connectivity structure as long as data are given with the calcium signal on a timescale of the order of 100 ms, because it cannot resolve the monosynaptic signal transmission of a few ms between neurons.

Here, we generated two-photon signals from the recorded spike trains by linearly convolving an exponential kernel function at each spike time. It has been reported that the fluorescence signals may exhibit nonlinearity with respect to the accumulated signals [42, 43]. Thus, it might be worthwhile to investigate whether the correlation between functional and monosynaptic connectivity could be strengthened by taking the nonlinearity into account.

While monosynaptic estimation is becoming more informative, particularly with the rapid development of high-density electrode techniques, functional connectivity analysis may also retain its importance because it may represent the influence of the invisible majority of neurons in the brain and also depend on behavioral contexts. The capabilities of two-photon recording technology seem to be developing even more rapidly, allowing the monitoring of a much larger number of neurons than microelectrode technology. Accordingly, an approach that integrates both monosynaptic and functional connectivity will influence future research design. This approach may reveal the conditions under which co-activation reflects underlying structure and how network dynamics modulate this relationship.

## METHODS

### An algorithm for estimating monosynaptic connections

Monosynaptic connectivity between a pair of neurons is detected by estimating the influence of the firing of one neuron on the firing of another neuron with a synaptic latency of a few ms. It has been proposed to construct a cross-correlation histogram or cross-correlogram and to determine the existence of a synaptic connection [12–14]. While the original theory took into account the statistical fluctuations in spike counts, *in vivo* neuronal firings show much larger fluctuations. There have been many efforts to resolve these confounding factors [15–29].

One of the authors and others have applied the Generalized Linear Model to absorb large fluctuations in cross-correlograms [24]. While this algorithm GLMCC drastically improved the estimation accuracy, we have newly found the cases where even this algorithm seems to make dubious inferences. We showed that this is caused by nondifferentiable fluctuations in the brain of awake animals, and revised the analysis algorithm to ShinGLMCC [29]. Given a set of recorded spike trains, ShinGLMCC provides a table of estimated connectivity for all neuron pairs, along with another table of their statistical significance *α* (Figure 1(a)). We have applied this method to the estimation of monosynaptic connections. The numerical codes of ShinGLMCC are available in a repository.

### Simulation of a large-scale network of neurons

We performed a numerical simulation of a network of neurons using the Multi-timescale Adaptive Threshold (MAT) model [38]. When we performed network simulations to test the ShinGLMCC in the reference [29], we chose interneuronal connections uniformly at random, taking into account neurophysiological knowledge such as balanced input, fractions of excitatory and inhibitory neurons, and lognormally distributed excitatory connectivity. In contrast, here we have adopted the connections estimated from the real data set. The peculiarity of the estimated connections is that they are clearly spatially localized, and that inhibitory neurons tend to have reciprocal feedback connections from excitatory neurons in each brain region (Figure 1).

The details of the simulation are as follows. The membrane potential *v* obeys a simple leaky integration of the input signal without reset: *τ*_*m*_*dv/dt* = −(*v* − *V*_L_) − (*α*_E_ + *β*_E_)(*v* − *V*_E_) − (*α*_*I*_ + *β*_*I*_)(*v* − *V*_*I*_), where *α*_E_ and *α*_I_ represent the relative excitatory and inhibitory conductances, respectively. The conductance evolves with 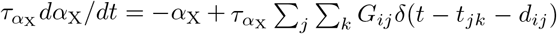 where X stands for excitatory E or inhibitory I, 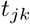 is the decay constant, *t*_*jk*_ is the *k*th spike time of the *j*th neuron, *d*_*ij*_ is a synaptic delay and *G*_*ij*_ is the synaptic weight from the *j*th neuron to the *i*th neuron. *δ*(*t*) is the Dirac delta function. *β*_X_ represents noise arising from background input, modeling the random bombardment from excitatory and inhibitory neurons in the outer population. This noise is described by a stochastic differential equation (Ornstein-Uhlenbeck process), with parameters selected such that its stationary distribution is a normal distribution with mean *β*_0,X_ and standard deviation *σ*_*β*,X_. The threshold *θ* (*t*) for spike generation is raised when the membrane potential is reached, and then subsequently decays with two timescales as *θ* (*t*) = ∑_*j*_ *H*(*t*−*t*_*j*_)+*ω, H*(*t*) = ∑_*k*=1,2_ *α*_*k*_ exp(−*t/τ*_*k*_), where *t*_*j*_ is the *j*th spike time of a neuron, *ω* is the resting value of the threshold, *τ*_*k*_ is the *k*th time constant, and *α*_*k*_ is the weight of the *k*th component (*k* = 1, 2). The shorter timescale *τ*_1_ = 10 ms represents the membrane time constant, and the longer timescale *τ*_2_ = 200 ms represents the adaptation of neuronal spiking.

In the present numerical simulation, we have assigned the excitatory or inhibitory nature of each neuron according to the dominance of excitatory or inhibitory output connections, and when the numbers were equal, we chose the type with the highest certainty that had the lowest *p*-value. We rescaled the amplitude of the background noise *β*_X_ for each neuron so that its simulated firing rate matched that observed in the recorded data. The other parameter values were adopted from those in the reference [29].

The connection matrix *{G*_*ij*_*}* for the network simulation was chosen to be proportional to the estimated connection matrix {*C*_*ij*_} : *G*_*ij*_ = *AC*_*ij*_ for all pairs of neurons *i* and *j*. The only one free parameter for this simulation is this coefficient *A*, which controls the connection strength between neurons. With a small *A*, the neurons will not show significant interdependence, while with a large *A*, the spike trains may show extra large fluctuations. We chose this parameter *A* so that the monosynaptic connections estimated from simulated spike trains are similar to those estimated between real spike trains in terms of the MCC for the estimated connections. The optimal coefficient was found to be *A* = 1.0. The numerical simulation was carried out for a time interval of 3,600 s with a time step of 0.0001s. The numerical codes for this simulation are also available in a repository.

### Stimulation to a part of a network

To see how functional connectivity is affected by external inputs, we performed network simulations by applying weak stimulation to a small group of neurons. This was done by increasing the excitatory noise to the neurons *β*_E_ by a factor of 1.1 for 1 second every 20 seconds. As a result, the firing rates of the stimulated neurons increased by about 1.5 times for that 1 second. The stimulated neurons were selected from the thalamus (from number 1 to 76) or from the superficial layer of the visual cortex (from number 195 to 235). By recording spikes from all 242 neurons for 3,600 s, we estimated monosynaptic and functional connections as in the unstimulated case.

## Code availability

All the numerical codes we have used for the analysis are available in the repository at https://github.com/yasuhirotsubo/neuroscience

## Biological data

We have analyzed publicly sourced data of spike trains recorded in parallel from the brain of a freely moving rat using a high-density electrode probe, Neuropixels [10]. We analyzed spike trains recorded with a single probe simulataneously from thalamus, hippocampus, and visual cortex. We used spike trains of 242 units according to their “good” reliability evaluation.

## Acknowledgements

We thank Hideaki Shimazaki for valuable comments on this manuscript. We also thank the research groups of the references [10, 11] who made their experimental data available to the public, especially Professors Nicholas A. Steinmetz, Matteo Carandini, and Timothy D. Harris for providing technical information related to the analysis of these data. We thank ChatGPT-4 for its contributions, which were selectively referenced during the refactoring of the Python code in our study. S.S. was supported by JSPS KAKENHI 22H05163. Y.T. was supported by JSPS KAKENHI 22K12186, 22H05511, and 24H01254.

## Author contributions

S.S. designed the study and drafted the manuscript. Y.T. analyzed experimental data, and performed numerical simulations.

